# Fatiguing Exercise Reduces Cellular Passive Young’s Modulus in Human Vastus Lateralis Muscle

**DOI:** 10.1101/2024.03.07.583989

**Authors:** Grace E. Privett, Austin W. Ricci, Larry L. David, Karen W. Needham, Yong How Tan, Karina H. Nakayama, Damien M. Callahan

## Abstract

Previous studies demonstrated that acute, exercise-induced fatigue transiently reduces whole-muscle stiffness. Because reduced muscle stiffness at fatigue may contribute to increased injury risk and impaired contractile performance, the present study seeks to elucidate potential intracellular mechanisms underlying these reductions. To that end, cellular passive Young’s Modulus was measured in single, permeabilized muscle fibers from healthy, recreationally-active males and females. Eight volunteers (4 male, 4 female) completed unilateral, repeated maximal voluntary knee extensions until fatigue, after which percutaneous needle biopsies were performed on the fatigued (F) and non-fatigued (NF) Vastus Lateralis muscles. Muscle samples were processed for mechanical assessment and separately for imaging and phosphoproteomics. Single fibers were passively (pCa 8.0), incrementally stretched to 156% of the initial sarcomere length to assess Young’s Modulus, calculated as the slope of the resulting stress-strain curve at short (strain = 1.00-1.24 %Lo) and long (strain = 1.32-1.56 %Lo) fiber lengths. Titin phosphorylation was assessed by liquid chromatography followed by high-resolution mass spectrometry (LC-MS). Passive modulus was significantly reduced by fatigue at short and long lengths in male, but not female, participants. Fatigue increased phosphorylation of four serine residues located within the elastic region of titin and reduced phosphorylation at one serine residue but did not impact active tension nor sarcomere ultrastructure. Collectively, these results suggest muscle fatigue reduces cellular passive modulus in young males, but not females, concurrent with altered titin phosphorylation. These results provide mechanistic insight contributing to the understanding of sex-based differences in soft tissue injury and falls risk.

**Key Points Summary:**

- Previous studies have shown that skeletal muscle stiffness is reduced following a single bout of fatiguing exercise.
- Lower muscle stiffness at fatigue may increase risk for soft-tissue injury, however, the underlying mechanisms of this change are unclear.
- Our findings show that fatiguing exercise reduces passive Young’s modulus in skeletal muscle cells from males but not females, suggesting that intracellular proteins contribute to reduced muscle stiffness with fatigue in a sex-dependent manner.
- The phosphorylation status of the intracellular protein titin is modified by fatiguing exercise in a way that may contribute to altered muscle stiffness after fatiguing exercise.
- These results provide important mechanistic insight that may help explain why biological sex impacts risk for soft tissue injury in with repeated or high intensity mechanical loading in athletes and falls risk in older adults.

**New and Noteworthy:** Muscle fatigue has previously been shown to reduce musculotendinous stiffness, but the underlying mechanisms remain unclear. Our study presents novel evidence of fatigue-induced reductions in passive cellular Young’s Modulus in skeletal muscle from males, but not females, in conjunction with fatigue-induced alterations in titin phosphorylation. Collectively, these results suggest that intracellular mechanisms including titin phosphorylation may contribute to altered skeletal muscle stiffness following fatiguing exercise, and that this response is mediated by biological sex.

## Introduction

Skeletal muscle stiffness is a critical characteristic of muscle function. Through influence on rate and efficiency of force transduction from muscle to the tendon (Wilson & Flanagan, 2008), joint range of motion (Wilson *et al*., 1991) and joint stability (Blackburn *et al*., 2011), muscle stiffness impacts locomotion and functionality in everyday tasks. Previous studies using shear wave elastography (Andonian *et al*., 2016; Siracusa *et al*., 2019; Chalchat *et al*., 2020) and B-mode ultrasound (Kubo *et al*., 2001) suggest that fatiguing exercise reduces whole skeletal muscle stiffness, which may be maladaptive. Given that fatigue also affects joint kinematics (Kernozek *et al*., 2008), and neuromuscular recruitment (Bouillard *et al*., 2014; De Ste Croix *et al*., 2015), fatigue-induced reduction of skeletal muscle stiffness may contribute to knee joint destabilization and subsequent soft tissue injury in athletes (Myer *et al*., 2008a; Watsford *et al*., 2010; Blackburn *et al*., 2011). Specifically, reduced muscle stiffness decreases the amount of strain elastic energy that can be absorbed by a tissue before injury occurs (Mair *et al*., 1996). When experienced in older adults, fatigue-induced muscle compliance may reduce joint stability and impair the ability to instantly adjust muscle function in response to perturbation, collectively contributing to increased falls risk at fatigue (Morrison *et al*., 2016). Despite the clear clinical implications of reduced musculotendinous stiffness following fatiguing exercise, the underlying mechanisms of this phenomenon, and whether they are mediated by biological sex, are not yet known.

Although previous measures of fatigue-induced reduction of whole-muscle stiffness included exclusively (Siracusa *et al*., 2019; Chalchat *et al*., 2020) or predominantly (Andonian *et al*., 2016) male participants, muscle stiffness appears to be influenced by biological sex, with women exhibiting reduced muscle stiffness compared to men (Kubo *et al*., 2003; Morse, 2011). These divergent mechanical properties have been attributed to the effects of sex hormones on whole-muscle (Chidi-Ogbolu & Baar, 2019; Ham *et al*., 2020) and tendon (Hansen & Kjaer, 2016) stiffness. In muscle, estrogen appears to support the maintenance of mass and strength, metabolic function, and connective tissue collagen turnover and incorporation (Chidi-Ogbolu & Baar, 2019). Estrogen also renders ligaments and tendons more compliant in females compared to males (Hansen & Kjaer, 2016). More compliant connective tissues increase joint laxity, which likely increases risk of anterior cruciate ligament (ACL) injury (Myer *et al*., 2008b). In fact, female athletes are more susceptible to knee injuries (Deitch *et al*., 2006), commonly to the ACL (Arendt *et al*., 1999; Matzkin & Garvey, 2019) than age-matched male athletes. By contrast, males experience greater inflammation following extreme muscle loading (Stupka *et al*., 2000) and male athletes experience greater incidence of muscle strains in practice and competition (Cross *et al*., 2013; Dalton *et al*., 2015). To complicate matters further, the effect of the menstrual cycle on skeletal muscle stiffness is unclear, with some studies demonstrating varied stiffness throughout the menstrual cycle (Ham *et al*., 2020) and others demonstrating no change (Bell *et al*., 2011). Oral contraceptive (OC) use appears to ameliorate cyclical estrogen fluctuations, and has therefore been studied as a possible approach to minimizing soft-tissue injury risk (Morse *et al*., 2013; Konopka *et al*., 2019). While some studies demonstrated altered muscle stiffness during OC use (Morse *et al*., 2013), others observed no change (Bell *et al*., 2011). Collectively, these studies suggest that biological sex does impact skeletal muscle stiffness, but the underlying mechanisms are poorly understood. Furthermore, whether biological sex mediates acute reduction of muscle stiffness following fatiguing exercise is also unclear.

Skeletal muscle stiffness is impacted by intracellular (sarcomeric proteins) and extracellular (extracellular matrix, ECM) elements, both of which can be altered. In cases of chronic effectors of stiffness, such as aging (Wood *et al*., 2014; Noonan *et al*., 2020b; Pavan *et al*., 2020) and physical exercise training (Noonan *et al*., 2020a), altered stiffness is commonly attributed to the remodeling of ECM collagen. Interestingly, observations of modified stiffness in chemically permeabilized single skeletal muscle fibers have been made in aged (Lim *et al*., 2019; Noonan *et al*., 2020b) and trained (Noonan *et al*., 2020a) muscle, suggesting that intracellular proteins also contribute to chronic stiffness adaptations. Within muscle fibers, passive stiffness is primarily determined by the visco-elastic protein titin (Ottenheijm *et al*., 2012; Lim *et al*., 2019). All three titin isoforms – two cardiac (N2B and N2BA) and one skeletal (N2A) – contain extensible (I-band) and non-extensible (A-band and M-band) regions. Titin isoform distribution can shift as a mechanism for tuning the stiffness of a muscle tissue. In cardiac muscle shifts occur between N2B and N2BA isoforms (Bupha-Intr *et al*., 2011), and in skeletal muscle the N2A isoforms vary in terms of titin size (Prado *et al*., 2005). Acutely, titin extensible regions are subject to post-translational modifications (PTMs), many of which have the potential to alter titin-based stiffness in response to stimuli such as fatiguing exercise.

Titin-based mechanisms of altered passive stiffness have been extensively studied in cardiac muscle, due to the clinical implications of altered cardiac titin for cardiomyopathies (Fukuda et al., 2005; Granzier & Irving, 1995; LeWinter & Granzier, 2014; Müller et al., 2014; Yamasaki et al., 2002). However, titin’s role in modulating skeletal muscle stiffness has been of growing interest, revealing altered titin-based stiffness in conditions such as cerebral palsy (Mathewson *et al*., 2014) and Ehler’s Danlos Syndrome (Ottenheijm *et al*., 2012). In both skeletal and cardiac muscle, the most studied titin PTM is phosphorylation. Phosphorylation sites have been identified in the immunoglobulin (Ig), PEVK, and unique sequence regions of the elastic I-band of titin, yet the locations of these phosphorylation sites vary across titin isoforms (Hamdani et al., 2017). One group (Müller *et al*., 2014) demonstrated that a single bout of exercise was sufficient to modify titin phosphorylation in association with altered myocyte stiffness in murine cardiac muscle. These results suggest that altered titin phosphorylation may contribute to acute, exercise-induced changes in muscle stiffness. Furthermore, such changes may have implications for whole-muscle stiffness, given titin’s influence on whole-muscle mechanical properties (Brynnel *et al*., 2018).

Despite growing support for a role of titin in regulating cardiac (Granzier & Irving, 1995; Yamasaki *et al*., 2002; Fukuda *et al*., 2005; Müller *et al*., 2014; LeWinter & Granzier, 2014) and skeletal (Ottenheijm *et al*., 2012; Mathewson *et al*., 2014; Müller *et al*., 2014; Brynnel *et al*., 2018) muscle stiffness, the contribution of acute titin phosphorylation to regulation of skeletal muscle stiffness in humans remains unclear. Therefore, the purpose of this study was to compare cellular passive stiffness, quantified as passive Young’s Modulus to account for potential variation in single fiber size, in skeletal muscle samples obtained from fatigued (F) and non-fatigued (NF) vastus lateralis muscles of healthy, young males and females. To interrogate potential mechanisms contributing to altered stiffness following fatigue, titin phosphorylation was compared in F versus NF skeletal muscle samples using liquid chromatography coupled with high resolution mass spectrometry (LC-MS). We hypothesized that fatiguing exercise would reduce passive modulus in single fibers from males and females in conjunction with increased titin phosphorylation.

## Methods

### Population

This protocol was approved by the Institutional Review Board at the University of Oregon. Eight young (18-21 yrs.) males (n=4) and females (n=4) from the University of Oregon and surrounding community consented to participate in this study. All participants were recreationally active yet reported no participation in structured physical exercise and no resistance training. Thus, they were characterized as “untrained”. Self-reported physical activity was confirmed by ActivePal (Pal Technologies, Glasgow, UK) accelerometers as described previously (Dowd *et al*., 2012). To limit the potential for menstrual cycle-dependent variation in circulating estradiol to contribute to variability in skeletal muscle mechanical properties, all female volunteers self-reporting eumenorrhea (n=3) were tested in the pre-follicular phase of the menstrual cycle (within 5 days of menses onset). The remaining female volunteer reported use of hormonal contraceptive and study timing was not considered with respect to menstrual cycle. Participants reported no orthopedic limitations (severe osteoarthritis, joint replacement or other orthopedic surgery in the previous six months), endocrine disease (hypo/hyper thyroidism, Addison’s Disease or Cushing’s syndrome), uncontrolled hypertension (>140/90 mmHg), neuromuscular disorder, significant heart, liver, kidney or respiratory disease, and/or diabetes. Participants were non-tobacco-smokers and had no current alcohol disorder. Finally, participants taking medications known to affect muscle stiffness or beta-adrenergic signaling of neuromuscular activation (including but not limited to beta blockers, calcium channel blockers, and muscle relaxers) or anabolic steroids were not included.

### Study Design

Participants visited the lab on 2 occasions separated by at least 1 week. During the first visit, volunteers habituated to measures of voluntary strength, power, and the fatigue of their dominant knee extensors (KE). During the second visit, volunteers performed measures of maximal voluntary isometric KE strength followed by fatiguing exercise to task failure. Fatiguing exercise was followed by bilateral, percutaneous needle muscle biopsies: one on the exercised limb immediately following exercise (“fatigued”) and the second on the contralateral, non-exercised limb (“non-fatigued”).

### Fatigue Protocol

Participants exercised the dominant limb on a Biodex System 3 dynamometer (Biodex Medical Systems, Shirley, NY). Participants were seated on the dynamometer with hips and knee flexed at 90° (180° = full extension). Prior to fatiguing exercise, participants completed three maximum voluntary isometric contractions (MVIC) of the knee extensors while analog voltage data for torque were sampled at 500 Hz. Analog data were converted to digital using an analog-to-digital converter (Cambridge Electronic Design, UK). Real-time visual feedback was provided to the participant to encourage maximal effort during the MVIC. The average torque value of the three MVICs was used to set the applied load to 30% MVIC maximal torque for the bout of fatiguing exercise. Following initial MVICs, participants performed repeated, voluntary knee extensions at a rate of 1 contraction per 1.5s at this isotonic load until task failure. Task failure was identified as the inability to perform knee extension through at least 50% of the range of motion. For seven of the eight participants, fatigue was quantified as the 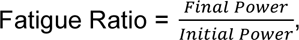 where “initial power” represents the average peak power of the first five knee extensions performed during fatiguing exercise, and “final power” represents the average peak power from the last five knee extensions. Due to a technical limitation, data were not appropriately saved to allow assessment of the fatigue ratio of one young female. Time to fatigue (task failure) was recorded for all eight participants.

### Muscle Biopsy Procedure

Percutaneous biopsy of the vastus lateralis (VL) muscle was performed under sterile conditions as previously detailed (Tarnopolsky *et al*., 2011). After sterilization of the biopsy site and injection of local anesthetic (1 or 2% lidocaine HCL [Hospira Worldwide, Lake Forest, IL, USA]), a small (∼5mm) incision was made in the skin and muscle fascia, allowing passage of a Bergstrom biopsy needle (5 mm diameter) to the belly of the VL muscle at a depth of ∼2-3 cm.

### Tissue Processing

VL biopsy typically yields between 100 and 150 mg (wet weight) of muscle tissue. After samples were removed from the biopsy needle, tissue was divided for proteomics (∼30 mg), single muscle fiber mechanics (∼40 mg), and electron microscopy (EM, ∼10 mg). Samples intended for proteomics analyses were immediately flash-frozen in liquid nitrogen and stored at -80°C. Sample intended for mechanical experimentation was placed in dissecting solution (MDS, 120.782 mM NaMS, 5.00 mM EGTA, 0.118 mM CaCl_2_, 1.00 mM MgCl_2_, 5.00 mM ATP-Na_2_H_2_, 0.25 mM KH_2_PO_4_, 20.00 mM BES, 1.789mM KOH) and parsed into bundles of ∼50 fibers. Bundles were tied to glass rods, chemically skinned overnight, and advanced through solutions of increasing glycerol content before long-term storage in 50% glycerol solution (5.00 mM EGTA, 2.50 mM MgCl_2_, 2.50 mM ATP-Na_2_H_2_, 10 mM imidazole, 170.00 mM potassium propionate, 1.00 mM sodium azide, 50% glycerol by volume) at -20°C. Sample apportioned for EM was tied to a glass rod, stretched slightly, and stored at 4°C in Karnovsky’s solution until embedding and sectioning as described elsewhere (Miller *et al*., 2009).

### Single fiber morphology and contractile measures

Prior to mechanical assays, fiber bundles and dissected single fibers were chemically skinned (MDS + 1% Triton X-100) before being transferred to plain MDS at ∼4°C until experimentation. Prepared fibers were mounted in relaxing solution (67.286 mM NaMS, 5.00 mM EGTA, 0.118 mM CaCl_2_, 6.867 mM MgCl_2_, 0.25 mM KH_2_PO_4_, 20.00 mM BES, 0.262 mM KOH, 1.00 mM DTT, 5.392 mM Mg-ATP, 15.00 mM CP, 300 U/mL CPK) between a force transducer and a length motor (Aurora Scientific, Inc., Aurora, ON, Canada) using the Moss clamp technique (Moss, 1979). Multiple wells are present under the mounting surface, allowing rapid transfer of the fiber between relaxing, pre-activating (81.181 mM NaMS, 5.00 mM EGTA, 0.012 mM CaCl_2_, 6.724 mM MgCl_2_, 5.00 mM KH_2_PO_4_, 20.00 mM BES, 1.00 mM DTT, 5.397 mM Mg-ATP, 15.00 mM CP, 300 U/mL CPK) and activating (57.549 mM NaMS, 5.00 mM EGTA, 5.021 mM CaCl_2_, 6.711 mM MgCl_2_, 5.00 mM KH_2_PO_4_, 20.00 mM BES, 9.674 mM KOH, 1.00 mM DTT, 5.437 mM Mg-ATP, 15.00 mM CP, 300 U/mL CPK) solutions. The mounting chamber contains a glass bottom and two prisms mounted to the side walls to allow for viewing of the fiber in top-down and side-view orientations. Fiber dimensions were measured as follows: d_top_ = average of three diameter measures along the fiber using the top-down view; d_side_ = average of three diameter measures along the fiber using the side view; fiber length = distance between the two trough edges (Callahan *et al*., 2015). Diameter measures were used for calculation of stress (force per unit area, kPa), and fiber length was used to calculate passive strain (change in length divided by original length, %L_0_). All mounted fibers were set to sarcomere length (SL) 2.65 µm. This SL is optimal for length-tension generation in human skeletal muscle (Burkholder & Lieber, 2001). Active tension was measured at SL 2.65 µm by moving the fiber to pre-activating solution followed by activating solution until a steady state tension was recorded. All fibers were activated (pCa 4.5) prior to passive stretching to measure active tension and confirm fiber viability.

### Passive stretch protocol

Passive modulus measures were performed in relaxing solution (pCa 8.0) using a passive stretch protocol adapted from previous work (Lim *et al*., 2019). Initial sarcomere length was set to 2.4 µm, followed by 7 incremental stretches to reach a final length of 156% of initial length (SL ∼ 3.7 µm). Each stretch lengthened the sample 8% of initial length and held this position for 2 minutes of stress-relaxation (Figure 1). Force response was measured by a force transducer and change in fiber length was measured by the displacement of the length motor. Sarcomere length was measured throughout the protocol using an inverted microscope located beneath the single fiber rig. To determine the extent to which acto-myosin interactions contributed to measures of passive modulus, a subset of fibers from two males was subjected to the passive stretch protocol in relaxing solution with the addition of 40 mM 2,3-butanedione monoxime (BDM), a myosin inhibitor. Following completion of the passive stretch protocol, each fiber was collected and placed in gel loading buffer (2% SDS, 62.5 mM Tris, 10% glycerol, 0.001% bromophenol blue, 5% β-mercaptoethanol, pH 6.8), centrifuged and heated at 65°C for 2 minutes, then stored at -80°C until later assessment of myosin heavy chain (MHC) isoform. Any fibers failing to demonstrate an increased force in response to fiber stretch (i.e. the subsequent force value after stress-relaxation was less than that of the previous stretch step) were excluded from analyses. The measured stress at the end of stress-relaxation was used for subsequent calculations and analyses. Data files were analyzed using custom code in Matlab software (R2020b, The MathWorks, Inc., Natick, Massachusetts).

**Figure 1.**
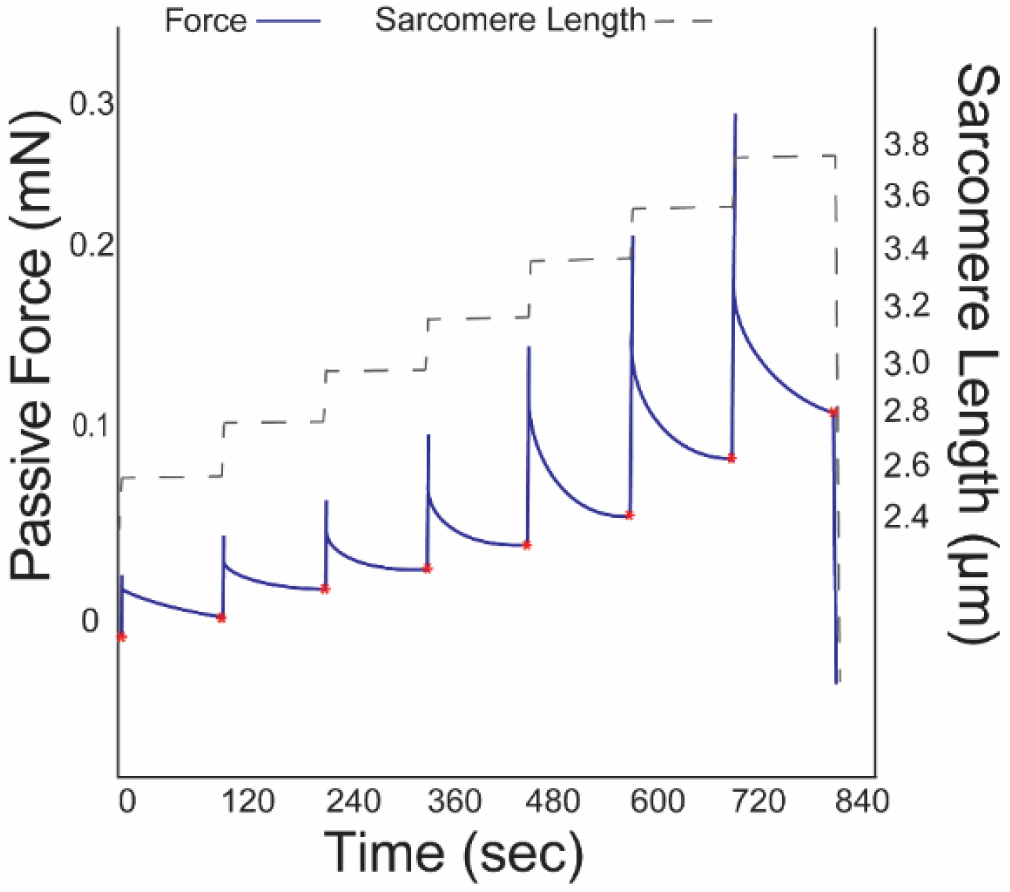
Sample force and sarcomere length traces produced during the passive stretch protocol. The force values measured at the end of each force decay, indicated by red asterisks, were used to calculate passive stress and, subsequently, passive Young’s Modulus.

### MHC isoform identification

Sodium dodecyl sulfate poly acrylamide gel electrophoresis (SDS-PAGE) was used to assess the MHC isoform of single muscle fibers. Sample from each fiber was loaded into its own well of a 4% stacking / 7% resolving polyacrylamide gel. The gel was run at 70 V for 3.5 hours followed by 200 V for 20 hours at 4°C (Miller *et al*., 2010). Gels were stained with a silver stain kit (Pierce Biotechnology, Waltham, MA) and the resulting MHC isoform (I, IIA, and/or IIX) expression was determined by comparison to a standard made from a multi-fiber homogenate (Figure 2).

**Figure 2.**
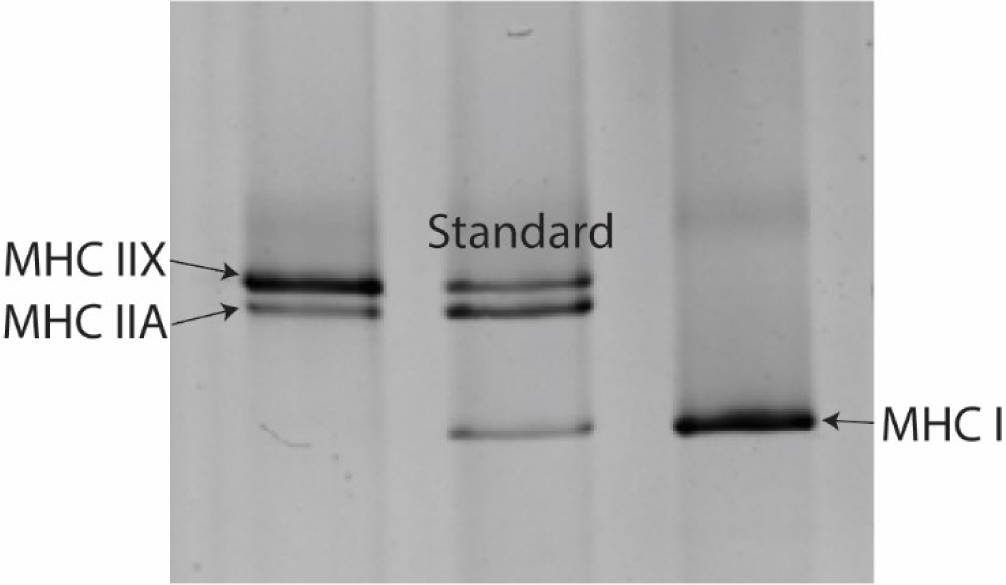
Sample image of silver-stained MHC bands. The left lane contains protein from a hybrid single fiber exhibiting both MHC IIA and MHC IIX isoforms. The middle band is a sample homogenate used to visualize all three MHC isoforms (I, IIA, IIX). The right lane contains protein from a single fiber expressing only MHC I isoform.

### Mass Spectrometry assessment of titin phosphorylation

Mass spectrometry was performed on skeletal muscle biopsy samples from individuals who were not included in the single fiber mechanics assays. This separate sample included four young (24.7 ± 5.7 years old), recreationally active individuals (2 males, 2 females) with a body mass index (BMI) of 24.4 ± 3.4 kg/m^2^. LC-MS was performed as previously described (Paulo *et al*., 2015) to determine which titin serine residues were differentially phosphorylated following acute fatigue. Briefly, samples were disrupted by shearing with glass beads, followed by protein digestion via trypsin and phosphopeptide purification by binding to TiO_2_ beads. Phosphopeptides were then labeled with one of 10 different tandem mass tags, combined into a single sample, and run via the Orbitrap Fusion mass spectrometer. Informatics methods (Plubell *et al*., 2017) were used to determine relative changes in phosphorylation of serine, threonine, and tyrosine residues.

### Electron Microscopy

Electron microscopy was performed to assess sarcomere ultrastructure (Miller *et al*., 2009). At the time of biopsy, samples apportioned for electron microscopy were fixed in Karnovsky’s Solution (2.5% glutaraldehyde, 2.5% formaldehyde, 0.1M Sodium Cacodylate). Before imaging, samples were treated with 2% osmium tetroxide in 0.5 M sodium cacodylate buffer (Ted Pella), stained with uranyl acetate, and embedded in epoxy resin. Cross sections (∼100 nm) were then cut using an ultramicrotome and contrasted with uranyl acetate before mounting on copper grids and subsequent imaging with a FEI Tecnai™ with iCorr™ Integrated Light and Transmission Electron Microscope. Qualitative assessment of sarcomere ultrastructure was used to check for disruption resulting from fatiguing exercise.

### Outcome Measures

Passive stress at each SL was calculated as 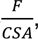 where “F” indicates the measured force value at the end of stress relaxation (Figure 1) and “CSA” indicates fiber elliptical cross-sectional area. 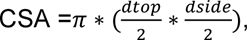 where “d_top_” is the average of three top diameter measures, and “d_side_” is the average of three side diameter measures. Strain was calculated as 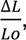 where “L_0_” indicates initial length. Passive stiffness was quantified as Young’s Modulus to account for potential differences in fiber size across samples. Passive Young’s Modulus was calculated as the slope of the stress-strain relationship at short fiber lengths (strain = 1.0-1.24 %Lo) and at long fiber lengths (strain = 1.32-1.56 %Lo). Separate slopes were calculated for short and long fiber lengths to consider the length dependence of cellular passive modulus measures (Noonan *et al*., 2020b). Additionally, the response of cellular passive modulus to fatiguing exercise was quantified for each participant by expressing the mean value for fatigued passive modulus as a percent of the mean value for non-fatigued modulus. Maximally activated tension was quantified as the measure of steady-state active force divided by fiber CSA. Titin phosphorylation, assessed via LC-MS, was quantified as fold change ratio from non-fatigued.

### Statistical Analyses

Statistical testing was conducted using SPSS software package (SPSS, IBM Corp., Armonk, NY, USA), unless otherwise specified. Anthropometric measures and activity data were compared between males and females using two-tailed, independent t-tests in excel. To evaluate differences in single fiber passive modulus at short and long lengths, a linear mixed model was run with fatigue, biological sex, and interaction terms as fixed effects and participant ID as a random effect to account for fiber variation within individuals, as described previously (Callahan *et al*., 2015). Subsequent analyses to investigate an interaction between biological sex and fatigue did so using separate linear mixed models in males and females, each including fatigue as a fixed effect and participant ID as a random effect. To test whether passive modulus was affected by treatment with BDM in a subset of fibers, a mixed effects model was run with fatigue condition and BDM treatment as main effects and participant ID as a random effect. To determine whether maximally activated tension, single fiber CSA, or fiber length differed by biological sex or fatigue condition, separate linear mixed effects models were run with sex, fatigue, and interaction terms as main effects and participant ID as a random effect. To assess titin phosphorylation of F versus NF sample using LC-MS, mean titin phosphorylation was quantified as fold change ratio, and any significant differences between F and NF samples were identified by a false detection rate (FDR) lower than 0.05. These LC-MS statistics were conducted using EdgeR software, proprietary to the Orbitrap Fusion.

### Data Availability Statement

Supporting data in results are presented in the manuscript and included in figures. Additional data are available as supporting information published with the paper on line.

## Results

### Descriptive Measures

The eight participants included were an average of 20.1 ± 1.1 years old. BMI (p=0.00878), height (p=0.0237), and weight (p=0.00665) were higher in males versus females (Table 1). Participants were not engaged in structured exercise training, and activity levels of individuals were confirmed by accelerometry. There were no significant differences in step count (p=0.646), minutes spent in light (< 75 steps/minute, p=0.137), moderate (75-125 steps/minute, p=0.424) or vigorous (>125 steps/minute, p=0.445) activity between males and females (Table 1). Despite efforts to select comparable numbers of MHC I and MHC II (including IIA, IIX, and A/X) fibers (Privett *et al*., 2024), MHC I fibers were under-represented in this sample (10% of total sample), especially in the male participants (3% of fibers from males). However, fibers expressing MHC IIA or MHC IIA/X were similarly represented in samples from males and females. Therefore, only fibers expressing MHC IIA and MHC IIA/X (n=129) were included in statistical analyses (Table 2). There was no significant difference in single fiber CSA between biological sexes (p=0.059) or fatigue conditions (p=0.566). Fiber length did not differ by biological sex (p=0.168) or fatigue condition (p=0.172).

**Table 1.** Anthropometric Data of Included Participants.

|  | N | BMI (kg/m<sup>2</sup>) * | Height (cm) * | Weight (kg) * | Step Count (steps/day) | Light Activity (minutes/day) | Moderate Activity (minutes/day) | Vigorous Activity (minutes/day) |
| --- | --- | --- | --- | --- | --- | --- | --- | --- |
| **Male** | 4 | $24.4 \pm 1.2$ | $185.3 \pm 13.0$ | $84.0 \pm 10.1$ | $7108 \pm 1525$ | $37.6 \pm 6.7$ | $50.8 \pm 14.7$ | $1.1 \pm 1.2$ |
| **Female** | 4 | $21.2 \pm 0.3$ | $160.3 \pm 5.5$ | $54.4 \pm 3.4$ | $7750 \pm 2174$ | $26.9 \pm 10.5$ | $60.7 \pm 18.0$ | $0.5 \pm 0.7$ |

\* Indicates significant difference between groups ( $p<0.05$ ). Data are shown as mean ± SD.

**Table 2.**
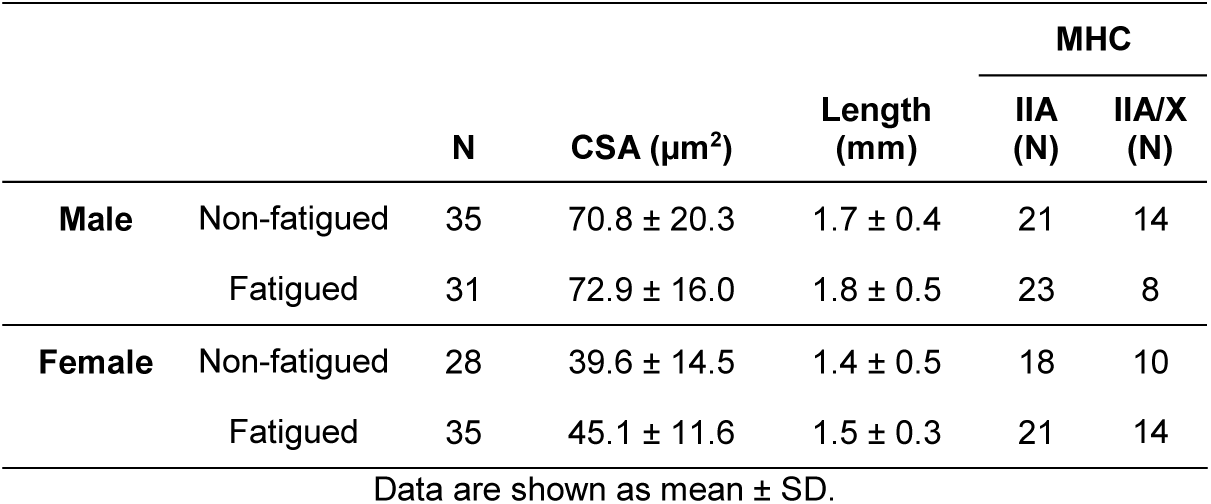
Summary Statistics of Fibers Analyzed.

### Fatiguing exercise

The average time to fatigue did not differ between males and females (74.6 ± 28.6 versus 69.4 ± 9.8 seconds, respectively, p=0.723). In the 4 males and 3 females for which data were collected, there was no difference in fatigue ratio by biological sex (0.41 ± 0.2 versus 0.33 ± 0.1, respectively, p=0.517).

### Assessment of passive stretch protocol and calculation of Young’s Modulus

To address potential concerns regarding incomplete stress decay after each applied strain, a 3^rd^ order polynomial function was used to perform a non-linear regression analysis, using the “fitnlm” function in MatLab, on the measured stress decay during the 112 seconds following applied strain. This produced a predicted final stress value that was compared to the measured stress at the end of each hold period. The difference between the measured stress and predicted stress, termed “missed stress”, was calculated at each stretch step, for each fiber. Given the greater stress decay expected at longer lengths (Figure 1), the missed stress values were assessed at strains 1.32-1.56% Lo. Stress decay was greatest at longer lengths, suggesting the greatest potential for incomplete stress decay. However, measured stress at longer lengths was only different from predicted stress by 0.024 ± 0.14%. Given this minimal variation, it is not likely that incomplete stress decay impacted the ability to draw meaningful conclusions from final measured stress, thus measured stress was used throughout. To assess the use of the dual-slope approach to calculating passive modulus, the coefficient of determination was calculated for raw data included in the short slope, the long slope, and the entire curve (both short and long slopes) for each fiber. The R^2^ values were not obviously different when calculated for the short slope (0.97 ± 0.00), long slope (0.97 ± 0.01), or the entire curve (0.95 ± 0.00). However, the use of the dual slope approach allowed for consideration of the sarcomere length-dependence of any significant effects observed (Noonan *et al*., 2020b). Specifically, the short slope covered the sarcomere length (SL) range = 2.4 ± 0.03 µm – 3.0 ± 0.10 µm and the long slope covered the SL range = 3.2 ± 0.13 µm – 3.8 ± 0.18 µm.

### Mechanical Measures

MHC isoform was determined by comparison of the band pattern of a single fiber to that of a multi-fiber homogenate, termed the “standard” (Figure 2) in a silver-stained acrylamide gel. Because MHC IIA and MHC IIA/X fibers comprised the majority of the sample, statistical analyses were only conducted on these fiber types. Passive modulus was not significantly different between males and females at short (p=0.353) nor long (p=0.978) fiber lengths. Although passive modulus was not significantly different in NF versus F fibers at short lengths (NF: 14.5 ± 5.6 kPa/%Lo, F: 14.1 ± 6.7 kPa/%Lo, p=0.341), passive modulus was significantly reduced in F fibers at long lengths (NF: 31.7 ± 8.6 kPa/%Lo, F: 28.6 ± 13.7 kPa/%Lo, p=0.043, Figure 3). Because the interaction of fatigue and biological sex was significant at both short (p=0.006) and long (p=0.020) lengths, the effect of fatigue on passive modulus was assessed separately in males and females (Figure 4A & B). Subsequent analyses revealed that fatigue-induced reductions in passive modulus were driven by males at short (NF: 14.6 ± 4.1 kPa/%Lo, F:11.6 ± 5.4 kPa/%Lo, p=0.002) and long (NF: 33.1±6.5 kPa/%Lo, F: 26.2 ± 8.8 kPa/%Lo, p<0.001) lengths, whereas modulus in single fibers from females was not significantly different with fatigue at short lengths (NF: 14.5 ± 7.0 kPa/%Lo, F: 16.2 ± 7.0 kPa/%Lo, p=0.263) or long lengths (NF: 30.0 ± 10.6 kPa/%Lo, F: 30.7 ± 16.7 kPa/%Lo, p=0.862, Figure 4C & D). Although males consistently demonstrated reduced passive cellular modulus in fatigued versus non-fatigued fibers at short (Figure 4E) and long (Figure 4F) lengths, the response in females varied considerably by individual, especially at long lengths. A separate set of fibers from two of the young males were treated with BDM and subjected to passive stiffness measures. In these samples, the effect of fatigue on cellular passive modulus was maintained at both short (p=0.027) and long (p=0.009) lengths (Figure 5), consistent with observations in the rest of the fibers from the male cohort. In short, BDM did not influence fatigue-induced reductions in cellular passive modulus. Considering cellular active contractile mechanics (Figure 6), there was no main effect of biological sex (p=0.189) or fatigue (p=0.603) on active isometric tension in this sample of MHC IIA and MHC IIA/X fibers.

**Figure 3.**
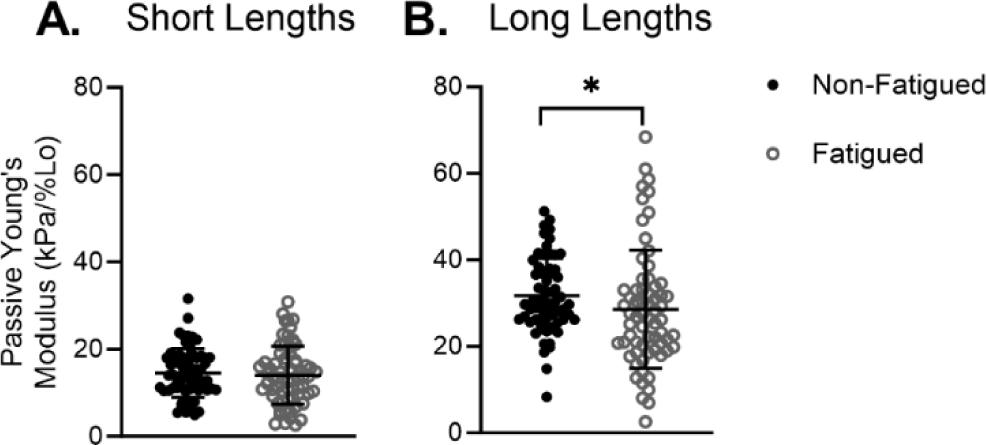
Passive modulus did not differ by fatigue condition at short lengths (A) but was significantly reduced in the fatigued sample at long lengths (B). * Indicates significant difference between fatigued and non-fatigued single fiber passive modulus (p<0.05). Data are shown as mean ± SD.

**Figure 4.**
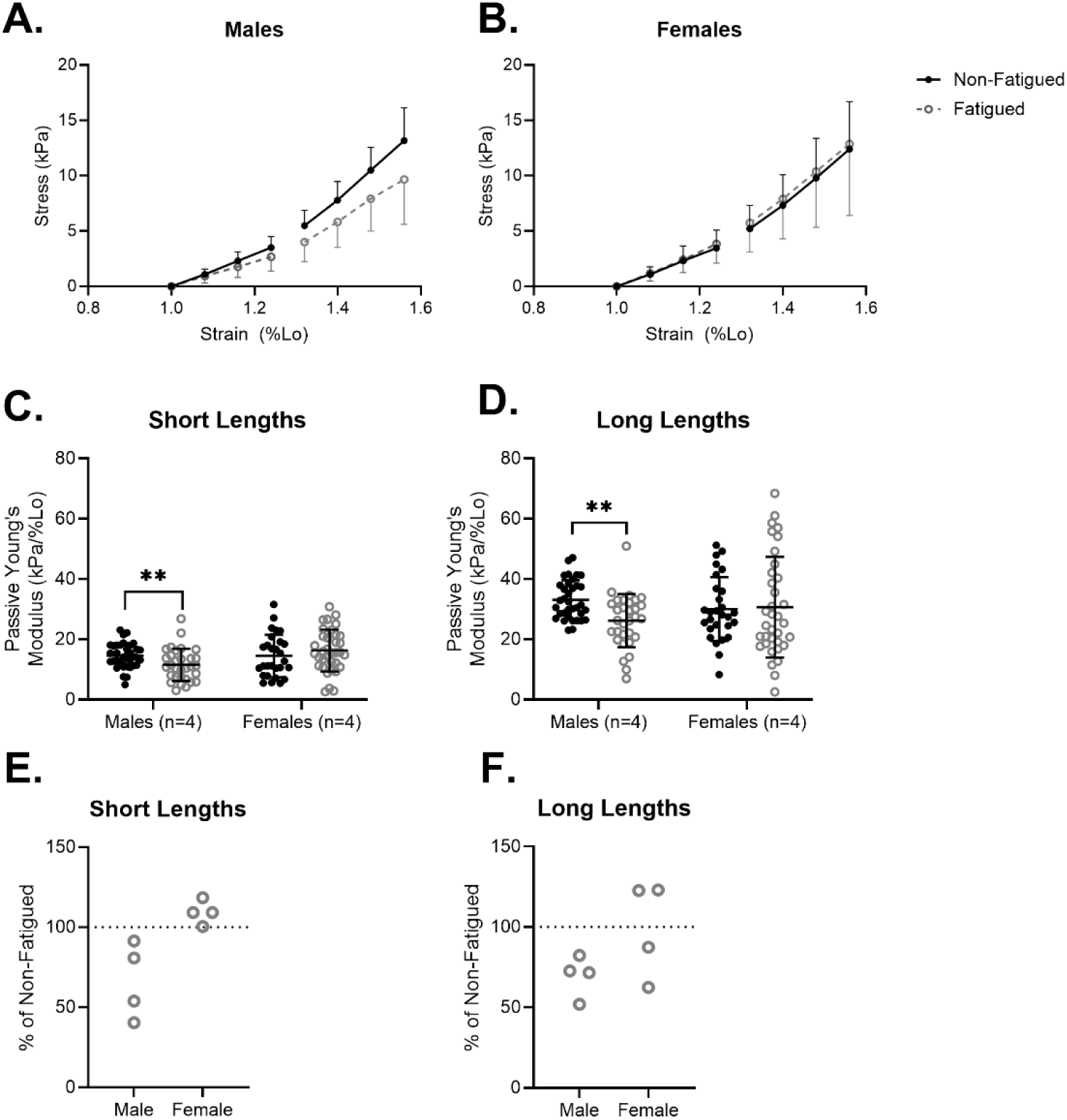
While there was an evident shift in slope of the stress-strain curve between non-fatigued and fatigued fibers at short and long lengths in the male group (A), there was no such shift in the female group (B). Statistical analysis confirmed that fatigue significantly reduced passive modulus in the single fibers of males at short (C) and long (D) fiber lengths but did not alter mean modulus in the fibers of females. Passive modulus of fatigued fibers was consistently lower compared to non-fatigued fibers in all four male participants at short (E) and long (F) lengths. However, females exhibited little to no change in passive modulus at short lengths, and variable responses at long lengths. Each individual point in figures 4E and 4F represents the average of all fibers, per individual participant. Data are shown as mean ± SD. ** indicates a significant fatigue effect (p<0.01).

**Figure 5.**
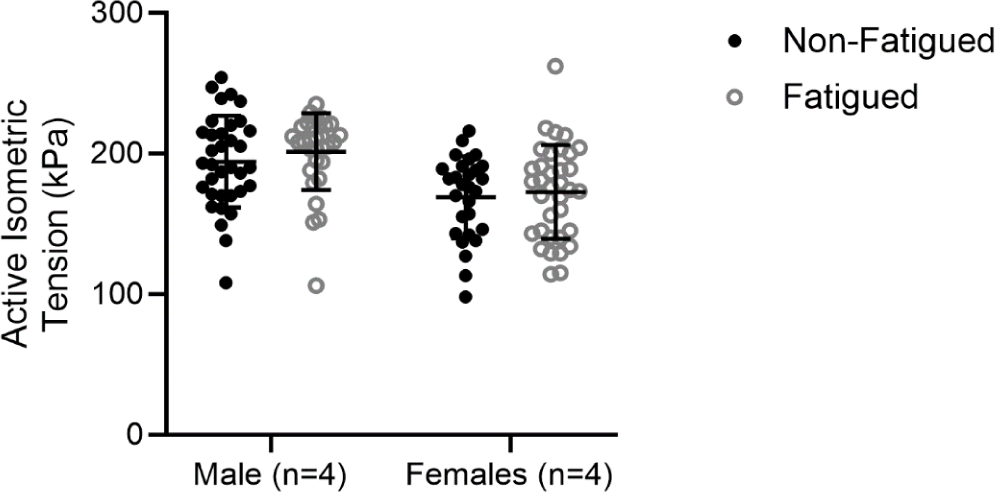
In a subset of fibers that were treated with BDM, passive modulus was significantly reduced in the F versus NF sample at both short and long fiber lengths, suggesting that fatigue-based differences in passive modulus persisted regardless of whether myosin was involved. Data are shown as mean ± SD. ** (p<0.01) and * (p<0.05) indicate significant differences by fatigue.

**Figure 6.**
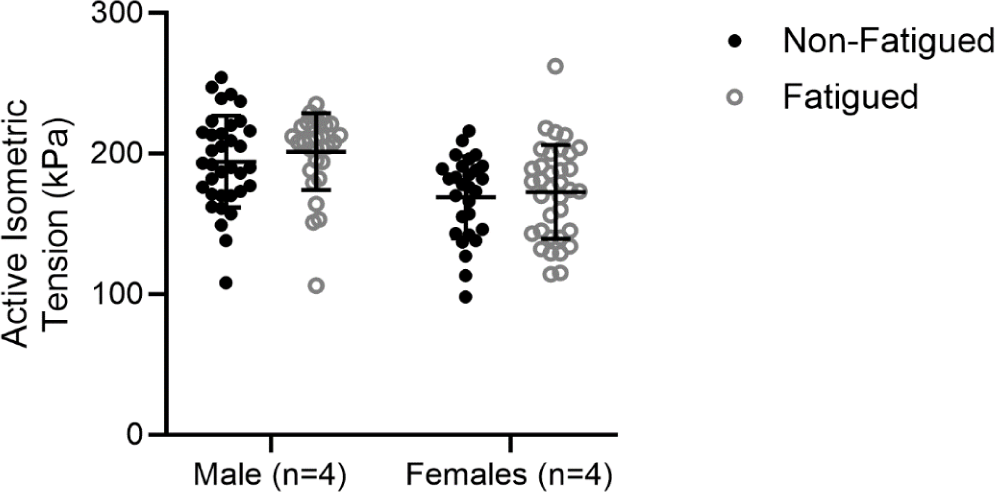
Cellular active tension was not significantly different between males and females (p=0.189) nor fatigue conditions (p=0.603). Data are shown as mean ± SD.

### Titin Phosphorylation

Mass spectrometry results (Figure 7) reveal increased phosphorylation of four titin serines: S12827 (FDR=0.0486), S12900 (FDR=0.0142), S12902 (FDR=0.0367), and S12918 (FDR=0.00195), and decreased phosphorylation at S28585 (FDR=0.0367). The Uniprot entry for human titin (entry Q8WZ42, The UniProt Consortium, 2023) suggests that S12827 is within immunoglobulin (Ig) domain 85 and S12900, S12902, and S12918 are located in between Ig domains 85 and 86. Uniprot also suggests that S28525 is located within Ig domain 132. A previous review summarizing Uniprot data (Hamdani *et al*., 2017) suggests that in human cardiac titin, S12827, S12900, S12902, and S12918 are located within the elastic I-band of titin and S28585 is located within the inelastic A band, yet it is unclear how similar serine locations are within titin substructures of cardiac versus skeletal titin isoforms.

**Figure 7.**
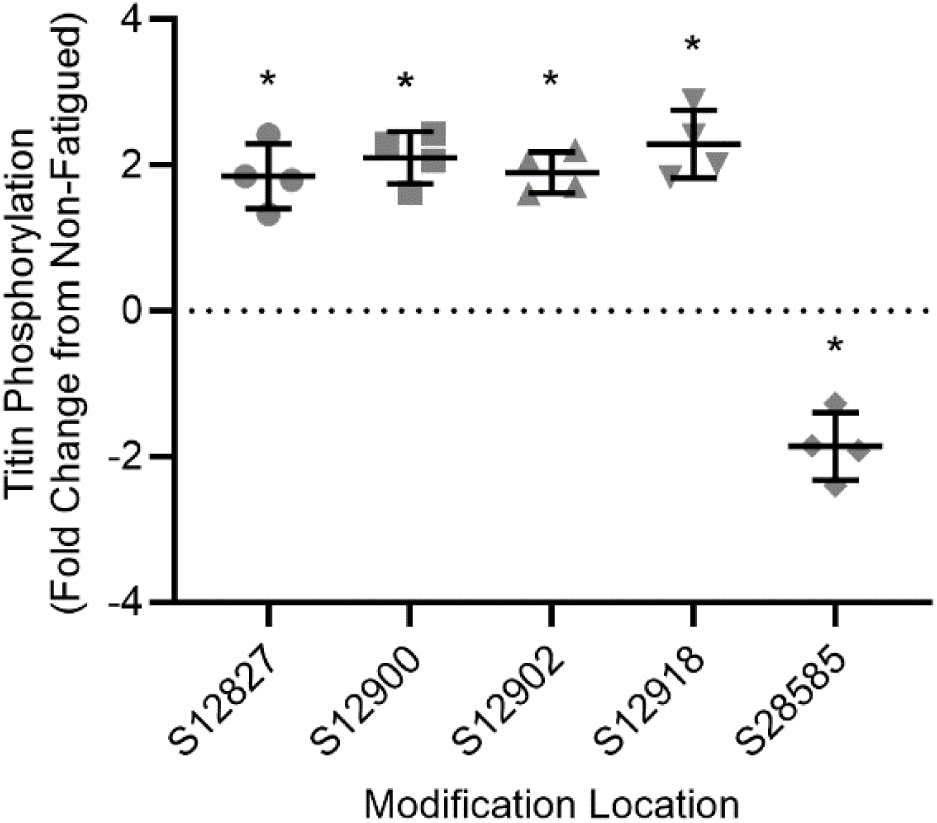
Mass spectrometry analysis demonstrated increased phosphorylation at 4 serine residues, S12827, S12900, S12902, and S12918 and decreased phosphorylation at serine residue S28525of the F versus NF sample. At each serine location, every point in the figure represents data form one participant (2 males, 2 females). Bars represent mean ± SD. * Indicates significantly different from NF (FDR<0.05).

### Electron Microscopy

Electron microscopy images do not demonstrate ultrastructural changes following fatiguing exercise (Figure 8). In contrast with published reports of contraction induced damage (Fridén et al., 1983; Fridén, 1984; Roth et al., 1999), which feature disordered sarcomeres and lack of registration between adjacent Z-lines, the sarcomeres visualized in this study were visibly similar between fatigued and non-fatigued samples.

**Figure 8:**
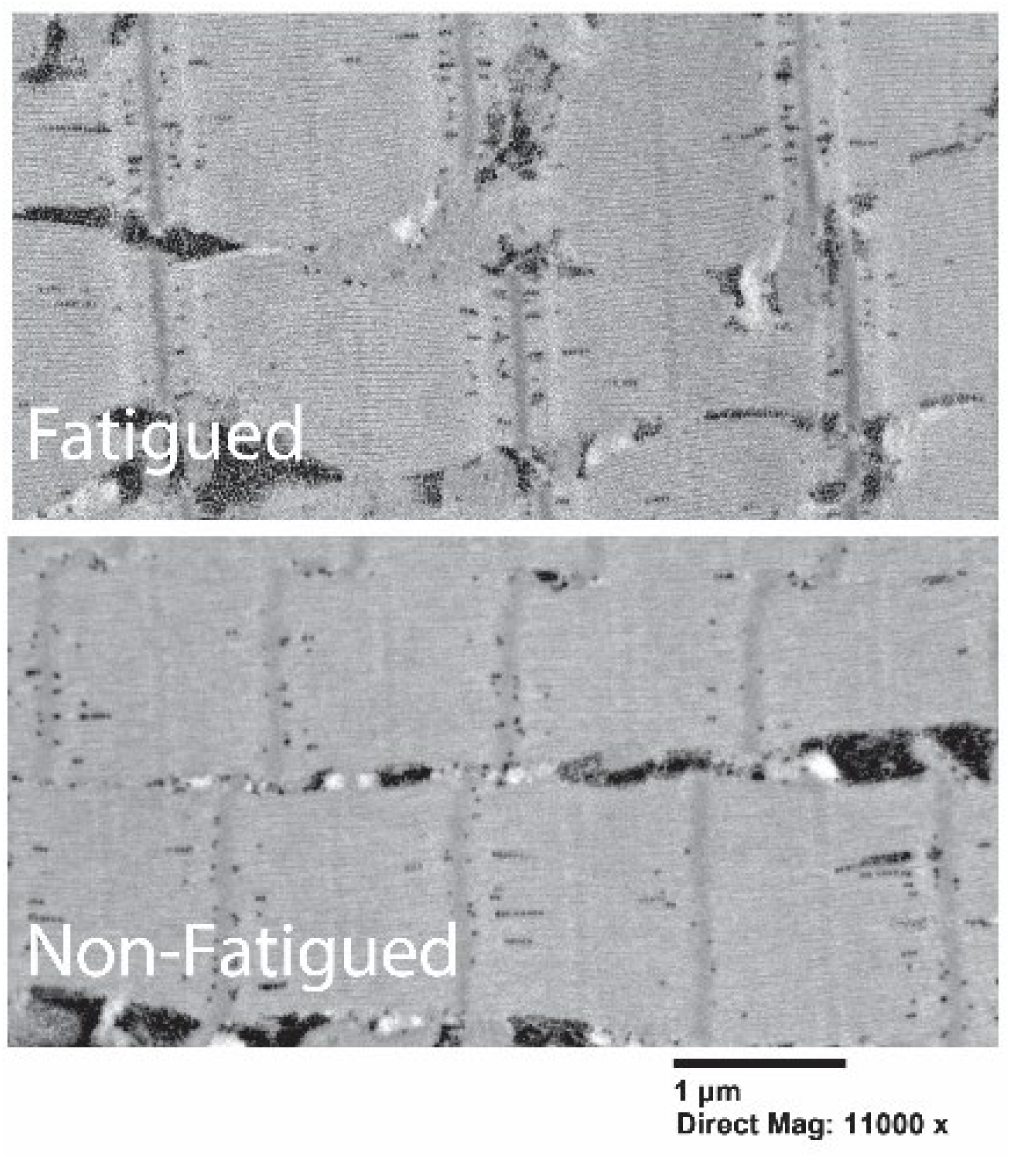
Representative electron microscopy images from fatigued and non-fatigued skeletal muscle samples did not show evidence of altered sarcomeric ultrastructure resulting from the fatigue protocol used in this study.

## Discussion

### Fatiguing exercise reduces cellular passive modulus

The present results demonstrate reduced passive modulus following fatiguing exercise in permeabilized single muscle fibers from young, untrained adults. Importantly, these observations were made in the absence of exercise-induced damage to the sarcomere ultrastructure (Figure 8). The observation of comparable active isometric tension in NF and F skeletal muscle fibers (Figure 6) further suggests that fiber integrity was not impaired in our fatigued sample. In these experiments, the use of chemical skinning on dissected single fibers, which permeabilizes the sarcolemma and removes ECM, supports the idea that reductions in modulus reflect changes to intracellular proteins, rather than to ECM mechanical properties. In particular, proteins likely to contribute to altered cellular passive modulus include contractile proteins actin and myosin and viscoelastic protein titin.

In a relaxing solution (pCa 8.0), the contribution of residual crossbridge formation to cellular passive modulus is possible. This idea was initially presented by D.K. Hill (Hill, 1968) and supported by later work suggesting that a small population of acto-myosin interactions contributed to force production in relaxed frog skeletal muscle (Campbell & Lakie, 1998). If a proportion of crossbridges do remain intact in relaxed muscle fibers, fatigue induced PTMs to myosin, presumably still present at the time of passive mechanics measurement, may affect changes in passive modulus via altered contribution of residual crossbridge formation. This possibility of residual crossbridge formation was considered in the present study. However, measurement of passive modulus in relaxing solution with 40 mM 2,3-butanedione monoxime (BDM), a myosin inhibitor, did not impact the observation of reduced passive modulus in F versus NF fibers (Figure 5), suggesting that residual crossbridge formation was not a primary contributor to fatigue-induced reduction of passive modulus. Instead, the reduced cellular passive modulus observed in this sample was more likely the result of a non-contractile intracellular mechanism.

At the cellular level, passive modulus is primarily determined by the sarcomeric protein titin (Lim et al., 2019; Ottenheijm et al., 2012) and there is evidence to support the role of titin in mediating whole-muscle passive stiffness (Brynnel et al., 2018). Titin-based stiffness can be modified by calcium (Nishikawa, 2020), heat shock proteins (HSPs, Kötter et al., 2014), oxidation (Alegre-Cebollada et al., 2014; Watanabe et al., 2020), and phosphorylation (Hamdani et al., 2017), all of which are upregulated during fatiguing exercise. Therefore, fatiguing exercise may modify titin-based stiffness through PTM of titin elastic domains. For example, exercise increases calcium cycling, prompting calcium-mediated binding of titin to the thin filament (Dutta *et al*., 2018) which shortens and stiffens the titin-based spring during active lengthening of skeletal muscle (Joumaa *et al*., 2008). Additionally, increased acidity during exercise promotes aggregation of titin Ig domains, which increases titin-based stiffness (Kötter *et al*., 2014).

However, previous work in human muscle fibers has demonstrated that HSPs also bind to titin Ig domains, thereby stabilizing unfolded domains and preventing aggregation of folded domains (Kötter *et al*., 2014). In this way, HSPs may prevent acidification-based increase of titin-based stiffness during exercise (Kötter *et al*., 2014). Exercise also induces oxidative stress, which results in numerous downstream mediators of titin-based stiffness (Beckendorf & Linke, 2015) including S-Glutathionylation, which has been shown to reduce stiffness of muscle cells (Alegre-Cebollada *et al*., 2014; Watanabe *et al*., 2020). In the present study, the storage solutions used included Dithiothreitol (DTT), an antioxidant, making it unlikely that titin oxidation contributed to the observed changes in cellular passive modulus.

Finally, exercise-induced β-adrenergic signaling, coupled with increased concentration of inorganic phosphate, likely contribute to titin phosphorylation (Krüger & Linke, 2006; Hamdani *et al*., 2017) during fatiguing exercise. Preclinical studies have demonstrated that binding of inorganic phosphate to titin can alter titin-based stiffness, though the nature of mechanical change is dependent on the location of phosphate binding (Müller *et al*., 2014). In the present study, five significant phosphorylation changes were identified in the fatigued sample of human skeletal muscle. Therefore, phosphorylation is a possible contributor to the mechanical changes observed following fatiguing exercise.

### Fatiguing exercise modifies titin phosphorylation in human skeletal muscle

Mass spectrometry analyses (Figure 7) suggest that the fatiguing exercise utilized in this study altered titin phosphorylation in human vastus lateralis muscle, and these phosphorylation events may have occurred in the elastic regions of titin. The notion that titin phosphorylation is one of the many PTMs with the capacity to alter titin-based stiffness is not new (Hamdani *et al*., 2017). In fact, it has been previously proposed that phosphorylation of titin Ig domains may reduce the ability of Ig domains to refold (Hamdani *et al*., 2017) which, given the importance of titin Ig refolding to elastic energy production (Rivas-Pardo *et al*., 2016), likely reduces titin-based stiffness. This proposed mechanism is similar to another mechanism described previously (Alegre-Cebollada *et al*., 2014; Watanabe *et al*., 2020), in which titin oxidation via S-glutathione inhibited titin Ig domain refolding and ultimately decreased the stiffness of human cardiomyocytes. Evidence of modified titin-based stiffness following increased or decreased titin phosphorylation is present in preclinical skeletal (Müller *et al*., 2014), preclinical cardiac (Fukuda *et al*., 2005), and human cardiac (Krüger *et al*., 2009) muscle research, and the specific stiffness response is evidently dependent on the location of the phosphorylation event. In the present study, five serine residues presumably located within the elastic regions of titin exhibited increased or decreased phosphorylation in F versus NF skeletal muscle, presenting the possibility that altered titin phosphorylation contributed to the changes observed in cellular passive modulus. The link between titin phosphorylation and cellular passive mechanics warrants further investigation as a potential mechanism underlying reduced musculoskeletal stiffness following fatiguing exercise.

In the present study, passive modulus was measured in a subset of single fibers treated with BDM to isolate the specific effects of titin on passive modulus. Furthermore, fatigue-induced reductions in passive modulus were greatest at the longest lengths observed (SL 3.2 ± 0.13µm – 3.8 ± 0.18µm), where the overlap of thick and thin filaments is minimal and passive modulus in permeabilized skeletal muscle cells is primarily titin-based. Nonetheless, it must be acknowledged that fatiguing exercise has the potential to modify sarcomeric proteins other than titin. Although titin is a likely contributor to the changes in passive mechanical properties observed, future studies in this area must keep in mind that contributions of other proteins are possible.

### The effect of fatiguing exercise on passive modulus is sex specific

The present results suggest that fatigue induces reduced skeletal muscle cellular passive modulus in a way that is specific to biological sex (Figure 4A & B). Specifically, cellular passive modulus was significantly reduced at short (Figure 4C) and long (Figure 4D) lengths in skeletal muscle cells from young, untrained males but not females. The lack of significance may be partially explained by the greater variety in response of passive modulus to fatiguing exercise in females compared to males, especially at longer lengths (Figure 4F). This variation in response might stem from the effects of sex hormones such as estrogen on skeletal muscle. Given the presence of estrogen receptors within skeletal muscle and the effects of estrogen on skeletal muscle strength (Collins *et al*., 2019), mitochondrial function (Pellegrino *et al*., 2022; Yoh *et al*., 2023), and regenerative capacity (Kitajima & Ono, 2016; Pellegrino *et al*., 2022), estrogen has a clear effect on skeletal muscle cellular function through modification of intracellular proteins. In fact, estradiol has been shown to directly affect regulatory light chain function in skeletal muscle (Lai *et al*., 2016), prompting the possibility that estrogen might also affect the function of other intracellular proteins, such as titin. In preclinical work using rat cardiac muscle, changes in circulating estrogen prompted shifts in titin isoform distribution in a way that altered cardiac stiffness (Bupha-Intr *et al*., 2011). This result suggests an influential role of estrogen on the regulation of titin-based stiffness. However, whether estrogen directly impacts titin-based stiffness in human skeletal muscle remains to be seen. Nonetheless, previous work has demonstrated a negative correlation between circulating estrogen and musculotendinous stiffness (Bell *et al*., 2012), supporting the notion that estrogen does impact whole skeletal muscle stiffness. Although, the effect of the menstrual cycle on whole-muscle stiffness remains unclear, with some evidence of stiffness variation throughout the menstrual cycle (Ham *et al*., 2020) and other evidence of no change (Bell *et al*., 2011). In males, previous work found no relationship between circulating estrogen and musculotendinous stiffness (Bell *et al*., 2012). Although the present study design attempted to control for fluctuation in circulating estrogen throughout the menstrual cycle by collecting biopsies at the same time point (pre-follicular phase) or in individuals using hormonal contraception, inter-individual differences in total circulating estrogen may still have been present in our female participants, perhaps contributing to the diversity in response of passive modulus to fatiguing exercise across female participants (Figure 4E and F). For this reason, future studies of cellular passive modulus in females should consider measuring circulating estrogen at the time of biopsy collection.

### Limitations

In this study, the uneven distribution of fiber-types limited our ability to test for a mediating effect of MHC on the response of cellular passive modulus to fatiguing exercise. Previous work (Miller *et al*., 2015) has suggested that cellular passive elastic modulus differs among fiber-types, highlighting the need to consider fiber-type in future studies of passive modulus. Importantly, the conclusions of the present study were not impacted by inclusion of all fiber-types (I, IIA, IIX, I/IIA, IIA/X) in the preliminary statistical analyses (data not shown). Finally, it must be acknowledged that although the location and exact sequence of titin serines are based on the highly annotated entry for human titin: Q8WZ42 (The UniProt Consortium, 2023), the precise phosphorylation locations detected in our mass spectrometry analyses, and where they fall along the titin protein, may not yet be fully understood. Therefore, our ability to speculate regarding potential functional implications of phosphorylation/de-phosphorylation at each titin serine is necessarily limited.

## Conclusions

In conclusion, this study presents novel evidence of reduced cellular passive Young’s Modulus following a single bout of fatiguing exercise. This observation parallels previous reports of reduced whole-muscle stiffness following fatiguing exercise (Andonian *et al*., 2016; Siracusa *et al*., 2019; Chalchat *et al*., 2020), supporting the notion that intracellular mechanisms contribute to acute reduction of whole-muscle stiffness. Furthermore, LC-MS data suggest that titin phosphorylation is altered at 5 serine residues during fatiguing exercise, which previous research (Müller *et al*., 2014) suggests may contribute to altered titin-based stiffness. Given prior evidence of fatigue-induced muscle compliance of whole-muscle was only recorded in males (Siracusa *et al*., 2019; Chalchat *et al*., 2020), it is perhaps not surprising that we only observed the effect at the cellular level in male participants. Perhaps more interesting, the clear link between muscle fatigue and injury risk (Mair *et al*., 1996) may also be related to sex-based differences in the dynamic response of muscle fiber compliance to fatigue. Indeed, while the biomechanical etiology of soft tissue injury risk is complicated, sex-dependent variation in fatigue-induced compliance provides a tantalizing link to sex-based variation in soft tissue injury. Our study attempted to limit variation in tissue mechanics in female participants by performing our measures at a time of presumably limited circulating estradiol. However, circulating estradiol was not directly measured and significant variation among and between female participants is quite possible. Perhaps this variation in circulating estradiol contributed to the interindividual variation in fatigue response in female participants (Figure 4E & F). Future studies of cellular passive modulus in females should consider whether circulating estrogen concentrations alter the response of cellular passive modulus to fatiguing exercise, as this might contribute to the increased incidence of soft-tissue injury in female athletes compared to their male counterparts (Arendt *et al*., 1999; Deitch *et al*., 2006; Matzkin & Garvey, 2019) or falls risk and resulting injury in older females (Stevens, 2005; Franse *et al*., 2017). Last, it remains to be seen whether the observations of altered passive modulus at the cellular level translate to the tissue level of skeletal muscle, an important step to understanding the interplay between intracellular and extracellular contributors to skeletal muscle stiffness and how they might be acutely altered by fatiguing exercise. Ultimately, the study of mechanisms underlying fatigue-induced reductions in skeletal muscle passive stiffness will contribute to efforts aimed at reducing fatigue-induced injury risk in athletes and falls risk in older adults.

## List of abbreviations

MHC: myosin heavy chain
SDS-PAGE: sodium dodecyl sulfate poly acrylamide gel electrophoresis
LC-MS: liquid chromatography followed by high resolution mass spectrometry
ECM: extracellular matrix
KE: knee extensors
MVIC: maximal voluntary isometric contraction
VL: vastus lateralis
SL: sarcomere length
PTM: post-translational modification
OC: oral contraceptive
ACL: anterior cruciate ligament
F: fatigued
NF: non-fatigued

## Author Contributions

These experiments were conducted in the Muscle Cellular Biology Laboratory in the University of Oregon Human Physiology Department. G.E.P., A.W.R., K.W.N, L.L.D., and Y.H.T. designed and performed experiments. G.E.P., A.W.R., K.W.N, L.L.D., Y.H.T., K.H.N., and D.M.C. analyzed and interpreted study data and revised the manuscript. G.E.P. drafted the manuscript. D.M.C. conceived of and directed the study. All authors approved the final version of the manuscript and agree to be accountable for all aspects of the work in ensuring that questions related to the accuracy or integrity of any part of the work are appropriately investigated and resolved. All persons designated as authors qualify for authorship and all those who qualify for authorship are listed.

## ACKNOWLEDGEMENTS

The Authors wish to thank the volunteers for their invaluable contributions to our work. Electron microscopy was performed at the Oregon Health & Science University Multiscale Microscopy Core. Proteomics assays were conducted by the Proteomics Shared Resource, directed by Dr. Ashok Reddy, at Oregon Health and Science University.

## REFERENCES

Alegre-Cebollada J, Kosuri P, Giganti D, Eckels E, Rivas-Pardo JA, Hamdani N, Warren CM, Solaro RJ, Linke WA & Fernández JM (2014). S-Glutathionylation of Cryptic Cysteines Enhances Titin Elasticity by Blocking Protein Folding. Cell 156, 1235–1246.

Andonian P, Viallon M, Le Goff C, de Bourguignon C, Tourel C, Morel J, Giardini G, Gergele L, Millet GP & Croisille P (2016). Shear-wave elastography assessments of quadriceps stiffness changes prior to, during and after prolonged exercise: a longitudinal study during an extreme mountain ultra-marathon. PLoS One 11, e0161855.

Arendt EA, Agel J & Dick R (1999). Anterior Cruciate Ligament Injury Patterns Among Collegiate Men and Women. J Athl Train 34, 86–92.

Beckendorf L & Linke WA (2015). Emerging importance of oxidative stress in regulating striated muscle elasticity. J Muscle Res Cell Motil 36, 25–36.

Bell DR, Blackburn JT, Norcorss MF, Ondrak KS, Hudson JD, Hackney AC & Padua DA (2012). Estrogen and muscle stiffness have a negative relationship in females. Knee Surg Sports Traumatol Arthrosc 20, 361–367.

Bell DR, Blackburn JT, Ondrak KS, Hackney AC, Hudson JD, Norcross MF & Padua DA (2011). The effects of oral contraceptive use on muscle stiffness across the menstrual cycle. Clinical Journal of Sport Medicine 21, 467–473.

Blackburn JT, Norcross MF & Padua DA (2011). Influences of hamstring stiffness and strength on anterior knee joint stability. Clinical Biomechanics 26, 278–283.

Bouillard K, Jubeau M, Nordez A & Hug F (2014). Effect of vastus lateralis fatigue on load sharing between quadriceps femoris muscles during isometric knee extensions. Journal of neurophysiology 111, 768–776.

Brynnel A, Hernandez Y, Kiss B, Lindqvist J, Adler M, Kolb J, Van der Pijl R, Gohlke J, Strom J & Smith J (2018). Downsizing the molecular spring of the giant protein titin reveals that skeletal muscle titin determines passive stiffness and drives longitudinal hypertrophy. Elife 7, e40532.

Bupha-Intr T, Oo YW & Wattanapermpool J (2011). Increased myocardial stiffness with maintenance of length-dependent calcium activation by female sex hormones in diabetic rats. American Journal of Physiology-Heart and Circulatory Physiology 300, H1661–H1668.

Burkholder TJ & Lieber RL (2001). Sarcomere length operating range of vertebrate muscles during movement. Journal of Experimental Biology 204, 1529–1536.

Callahan DM, Tourville TW, Miller MS, Hackett SB, Sharma H, Cruickshank NC, Slauterbeck JR, Savage PD, Ades PA & Maughan DW (2015). Chronic disuse and skeletal muscle structure in older adults: sex-specific differences and relationships to contractile function. American Journal of Physiology-Cell Physiology 308, C932–C943.

Chalchat E, Gennisson J-L, Peñailillo L, Oger M, Malgoyre A, Charlot K, Bourrilhon C, Siracusa J & Garcia-Vicencio S (2020). Changes in the viscoelastic properties of the vastus lateralis muscle with fatigue. Frontiers in Physiology 11, 307.

Chidi-Ogbolu N & Baar K (2019). Effect of Estrogen on Musculoskeletal Performance and Injury Risk. Frontiers in Physiology. Available at: https://www.frontiersin.org/article/10.3389/fphys.2018.01834 [Accessed May 5, 2022].

Collins BC, Laakkonen EK & Lowe DA (2019). Aging of the Musculoskeletal System: How the Loss of Estrogen Impacts Muscle Strength. Bone 123, 137–144.

Cross KM, Gurka KK, Saliba S, Conaway M & Hertel J (2013). Comparison of hamstring strain injury rates between male and female intercollegiate soccer athletes. Am J Sports Med 41, 742–748.

Dalton SL, Kerr ZY & Dompier TP (2015). Epidemiology of Hamstring Strains in 25 NCAA Sports in the 2009-2010 to 2013-2014 Academic Years. Am J Sports Med 43, 2671–2679.

De Ste Croix MBA, Priestley AM, Lloyd RS & Oliver JL (2015). ACL injury risk in elite female youth soccer: Changes in neuromuscular control of the knee following soccer-specific fatigue. Scandinavian Journal of Medicine & Science in Sports 25, e531–e538.

Deitch JR, Starkey C, Walters SL & Moseley JB (2006). Injury Risk in Professional Basketball Players: A Comparison of Women’s National Basketball Association and National Basketball Association Athletes. Am J Sports Med 34, 1077–1083.

Dowd KP, Harrington DM & Donnelly AE (2012). Criterion and Concurrent Validity of the activPAL^TM^ Professional Physical Activity Monitor in Adolescent Females. PLOS ONE 7, e47633.

Dutta S, Tsiros C, Sundar SL, Athar H, Moore J, Nelson B, Gage MJ & Nishikawa K (2018). Calcium increases titin N2A binding to F-actin and regulated thin filaments. Sci Rep 8, 14575.

Franse CB, Rietjens JA, Burdorf A, Grieken A van, Korfage IJ, Heide A van der, Raso FM, Beeck E van & Raat H (2017). A prospective study on the variation in falling and fall risk among community-dwelling older citizens in 12 European countries. BMJ Open 7, e015827.

Fukuda N, Wu Y, Nair P & Granzier HL (2005). Phosphorylation of titin modulates passive stiffness of cardiac muscle in a titin isoform-dependent manner. The Journal of general physiology 125, 257–271.

Granzier HL & Irving TC (1995). Passive tension in cardiac muscle: contribution of collagen, titin, microtubules, and intermediate filaments. Biophys J 68, 1027–1044.

Ham S, Kim S, Choi H, Lee Y & Lee H (2020). Greater muscle stiffness during contraction at menstruation as measured by shear-wave elastography. The Tohoku Journal of Experimental Medicine 250, 207–213.

Hamdani N, Herwig M & Linke WA (2017). Tampering with springs: phosphorylation of titin affecting the mechanical function of cardiomyocytes. Biophysical reviews 9, 225–237.

Hansen M & Kjaer M (2016). Sex hormones and tendon. Metabolic influences on risk for tendon disorders139–149.

Joumaa V, Rassier DE, Leonard TR & Herzog W (2008). The origin of passive force enhancement in skeletal muscle. American Journal of Physiology-Cell Physiology 294, C74–C78.

Kernozek TW, Torry MR & Iwasaki M (2008). Gender differences in lower extremity landing mechanics caused by neuromuscular fatigue. The American journal of sports medicine 36, 554–565.

Kitajima Y & Ono Y (2016). Estrogens maintain skeletal muscle and satellite cell functions. J Endocrinol 229, 267–275.

Konopka JA, Hsue LJ & Dragoo JL (2019). Effect of Oral Contraceptives on Soft Tissue Injury Risk, Soft Tissue Laxity, and Muscle Strength: A Systematic Review of the Literature. Orthopaedic Journal of Sports Medicine 7, 2325967119831061.

Kötter S, Unger A, Hamdani N, Lang P, Vorgerd M, Nagel-Steger L & Linke WA (2014). Human myocytes are protected from titin aggregation-induced stiffening by small heat shock proteins. Journal of Cell Biology 204, 187–202.

Krüger M, Kötter S, Grützner A, Lang P, Andresen C, Redfield MM, Butt E, Dos Remedios CG & Linke WA (2009). Protein kinase G modulates human myocardial passive stiffness by phosphorylation of the titin springs. Circulation research 104, 87–94.

Krüger M & Linke WA (2006). Protein kinase-A phosphorylates titin in human heart muscle and reduces myofibrillar passive tension. Journal of Muscle Research & Cell Motility 27, 435–444.

Kubo K, Kanehisa H & Fukunaga T (2003). Gender differences in the viscoelastic properties of tendon structures. European journal of applied physiology 88, 520–526.

Kubo K, Kanehisa H, Kawakami Y & Fukunaga T (2001). Effects of repeated muscle contractions on the tendon structures in humans. European journal of applied physiology 84, 162–166.

Lai S, Collins BC, Colson BA, Kararigas G & Lowe DA (2016). Estradiol modulates myosin regulatory light chain phosphorylation and contractility in skeletal muscle of female mice. American Journal of Physiology-Endocrinology and Metabolism 310, E724–E733.

LeWinter MM & Granzier HL (2014). Cardiac Titin and Heart Disease. J Cardiovasc Pharmacol 63, 207– 212.

Lim J-Y, Choi SJ, Widrick JJ, Phillips EM & Frontera WR (2019). Passive force and viscoelastic properties of single fibers in human aging muscles. European journal of applied physiology 119, 2339–2348.

Mair SD, Seaber AV, Glisson RR & Garrett WE (1996). The Role of Fatigue in Susceptibility to Acute Muscle Strain Injury. Am J Sports Med 24, 137–143.

Mathewson MA, Chambers HG, Girard PJ, Tenenhaus M, Schwartz AK & Lieber RL (2014). Stiff muscle fibers in calf muscles of patients with cerebral palsy lead to high passive muscle stiffness. Journal of Orthopaedic Research 32, 1667–1674.

Matzkin E & Garvey K (2019). Sex Differences in Common Sports-Related Injuries. NASN School Nurse 34, 266–269.

Miller MS, Bedrin NG, Ades PA, Palmer BM & Toth MJ (2015). Molecular determinants of force production in human skeletal muscle fibers: effects of myosin isoform expression and cross-sectional area. American Journal of Physiology-Cell Physiology 308, C473–C484.

Miller MS, VanBuren P, LeWinter MM, Braddock JM, Ades PA, Maughan DW, Palmer BM & Toth MJ (2010). Chronic heart failure decreases cross-bridge kinetics in single skeletal muscle fibres from humans. The Journal of Physiology 588, 4039–4053.

Miller MS, VanBuren P, LeWinter MM, Lecker SH, Selby DE, Palmer BM, Maughan DW, Ades PA & Toth MJ (2009). Mechanisms underlying skeletal muscle weakness in human heart failure: alterations in single fiber myosin protein content and function. Circulation: Heart Failure 2, 700–706.

Morrison S, Colberg SR, Parson HK, Neumann S, Handel R, Vinik EJ, Paulson J & Vinik AI (2016). Walking-induced fatigue leads to increased falls risk in older adults. Journal of the American Medical Directors Association 17, 402–409.

Morse C, Spencer J, Hussain A & Onambélé-Pearson G (2013). The effect of the oral contraceptive pill on the passive stiffness of the human gastrocnemius muscle in vivo. Journal of musculoskeletal & neuronal interactions 13, 97–104.

Morse CI (2011). Gender differences in the passive stiffness of the human gastrocnemius muscle during stretch. European journal of applied physiology 111, 2149–2154.

Moss RL (1979). Sarcomere length-tension relations of frog skinned muscle fibres during calcium activation at short lengths. The Journal of physiology 292, 177–192.

Müller AE, Kreiner M, Kötter S, Lassak P, Bloch W, Suhr F & Krüger M (2014). Acute exercise modifies titin phosphorylation and increases cardiac myofilament stiffness. Frontiers in physiology 5, 449.

Myer GD, Ford KR, Paterno MV, Nick TG & Hewett TE (2008a). The Effects of Generalized Joint Laxity on Risk of Anterior Cruciate Ligament Injury in Young Female Athletes. Am J Sports Med 36, 1073– 1080.

Myer GD, Ford KR, Paterno MV, Nick TG & Hewett TE (2008b). The effects of generalized joint laxity on risk of anterior cruciate ligament injury in young female athletes. The American journal of sports medicine 36, 1073–1080.

Noonan AM, Mashouri P, Chen J, Power GA & Brown SH (2020a). Training Induced Changes to Skeletal Muscle Passive Properties Are Evident in Both Single Fibers and Fiber Bundles in the Rat Hindlimb. Frontiers in Physiology 11, 907.

Noonan AM, Mazara N, Zwambag DP, Weersink E, Power GA & Brown SH (2020b). Age-related changes in human single muscle fibre passive elastic properties are sarcomere length dependent. Experimental Gerontology 137, 110968.

Ottenheijm CA, Voermans NC, Hudson BD, Irving T, Stienen GJ, Van Engelen BG & Granzier H (2012). Titin-based stiffening of muscle fibers in Ehlers-Danlos Syndrome. Journal of Applied Physiology 112, 1157–1165.

Paulo JA, McAllister FE, Everley RA, Beausoleil SA, Banks AS & Gygi SP (2015). Effects of MEK inhibitors GSK1120212 and PD0325901 in vivo using 10-plex quantitative proteomics and phosphoproteomics. Proteomics 15, 462–473.

Pavan P, Monti E, Bondí M, Fan C, Stecco C, Narici M, Reggiani C & Marcucci L (2020). Alterations of Extracellular Matrix Mechanical Properties Contribute to Age-Related Functional Impairment of Human Skeletal Muscles. International Journal of Molecular Sciences 21, 3992.

Pellegrino A, Tiidus PM & Vandenboom R (2022). Mechanisms of Estrogen Influence on Skeletal Muscle: Mass, Regeneration, and Mitochondrial Function. Sports Med 52, 2853–2869.

Plubell DL, Wilmarth PA, Zhao Y, Fenton AM, Minnier J, Reddy AP, Klimek J, Yang X, David LL & Pamir N (2017). Extended multiplexing of tandem mass tags (TMT) labeling reveals age and high fat diet specific proteome changes in mouse epididymal adipose tissue. Molecular & Cellular Proteomics 16, 873–890.

Prado LG, Makarenko I, Andresen C, Krüger M, Opitz CA & Linke WA (2005). Isoform diversity of giant proteins in relation to passive and active contractile properties of rabbit skeletal muscles. The Journal of general physiology 126, 461–480.

Privett GE, Ricci AW, Ortiz-Delatorre J & Callahan DM (2024). Predicting Myosin Heavy Chain Isoform from Post-Dissection Fiber Length in Human Skeletal Muscle Fibers. American Journal of Physiology-Cell Physiology; DOI: 10.1152/ajpcell.00700.2023.

Rivas-Pardo JA, Eckels EC, Popa I, Kosuri P, Linke WA & Fernández JM (2016). Work Done by Titin Protein Folding Assists Muscle Contraction. Cell Reports 14, 1339–1347.

Siracusa J, Charlot K, Malgoyre A, Conort S, Tardo-Dino P-E, Bourrilhon C & Garcia-Vicencio S (2019). Resting muscle shear modulus measured with ultrasound shear-wave elastography as an alternative tool to assess muscle fatigue in humans. Frontiers in physiology 10, 626.

Stevens JA (2005). Gender differences for non-fatal unintentional fall related injuries among older adults. Injury Prevention 11, 115–119.

Stupka N, Lowther S, Chorneyko K, Bourgeois JM, Hogben C & Tarnopolsky MA (2000). Gender differences in muscle inflammation after eccentric exercise. Journal of Applied Physiology 89, 2325–2332.

Tarnopolsky MA, Pearce E, Smith K & Lach B (2011). Suction-modified Bergström muscle biopsy technique: Experience with 13,500 procedures. Muscle & nerve 43, 716–725.

The UniProt Consortium (2023). UniProt: the Universal Protein Knowledgebase in 2023. Nucleic Acids Research 51, D523–D531.

Watanabe D, Lamboley CR & Lamb GD (2020). Effects of S-glutathionylation on the passive force–length relationship in skeletal muscle fibres of rats and humans. Journal of Muscle Research and Cell Motility 41, 239–250.

Watsford ML, Murphy AJ, McLachlan KA, Bryant AL, Cameron ML, Crossley KM & Makdissi M (2010). A Prospective Study of the Relationship between Lower Body Stiffness and Hamstring Injury in Professional Australian Rules Footballers. Am J Sports Med 38, 2058–2064.

Wilson GJ, Wood GA & Elliott BC (1991). The relationship between stiffness of the musculature and static flexibility: an alternative explanation for the occurrence of muscular injury. International journal of sports medicine 12, 403–407.

Wilson JM & Flanagan EP (2008). The Role of Elastic Energy in Activities with High Force and Power Requirements: A Brief Review. The Journal of Strength & Conditioning Research 22, 1705–1715.

Wood LK, Kayupov E, Gumucio JP, Mendias CL, Claflin DR & Brooks SV (2014). Intrinsic stiffness of extracellular matrix increases with age in skeletal muscles of mice. Journal of applied physiology 117, 363–369.

Yamasaki R, Wu Y, McNabb M, Greaser M, Labeit S & Granzier H (2002). Protein Kinase A Phosphorylates Titin’s Cardiac-Specific N2B Domain and Reduces Passive Tension in Rat Cardiac Myocytes. Circulation Research 90, 1181–1188.

Yoh K, Ikeda K, Horie K & Inoue S (2023). Roles of estrogen, estrogen receptors, and estrogen-related receptors in skeletal muscle: regulation of mitochondrial function. International Journal of Molecular Sciences 24, 1853.

